# Exploring putative enteric methanogenesis inhibitors using molecular simulations and a graph neural network

**DOI:** 10.1101/2024.09.16.613350

**Authors:** Randy Aryee, Noor S. Mohammed, Supantha Dey, B. Arunraj, Swathi Nadendla, Karuna Anna Sajeevan, Matthew R. Beck, A. Nathan Frazier, Jacek A. Koziel, Thomas J. Mansell, Ratul Chowdhury

## Abstract

Atmospheric methane (CH_4_) acts as a key contributor to global warming. As CH_4_ is a short-lived climate forcer (12 years atmospheric lifespan), its mitigation represents the most promising means to address climate change in the short term. Enteric CH_4_ (the biosynthesized CH_4_ from the rumen of ruminants) represents 5.1% of total global greenhouse gas (GHG) emissions, 23% of emissions from agriculture, and 27.2% of global CH_4_ emissions. Therefore, it is imperative to investigate methanogenesis inhibitors and their underlying modes of action. We hereby elucidate the detailed biophysical and thermodynamic interplay between anti-methanogenic molecules and cofactor F_430_ of methyl coenzyme M reductase and interpret the stoichiometric ratios and binding affinities of sixteen inhibitor molecules. We leverage this as prior in a graph neural network to first functionally cluster these sixteen known inhibitors among ∼54,000 bovine metabolites. We subsequently demonstrate a protocol to identify precursors to and putative inhibitors for methanogenesis, based on Tanimoto chemical similarity and membrane permeability predictions. This work lays the foundation for computational and *de novo* design of inhibitor molecules that retain/ reject one or more biochemical properties of known inhibitors discussed in this study.

**COVER ABSTRACT:** 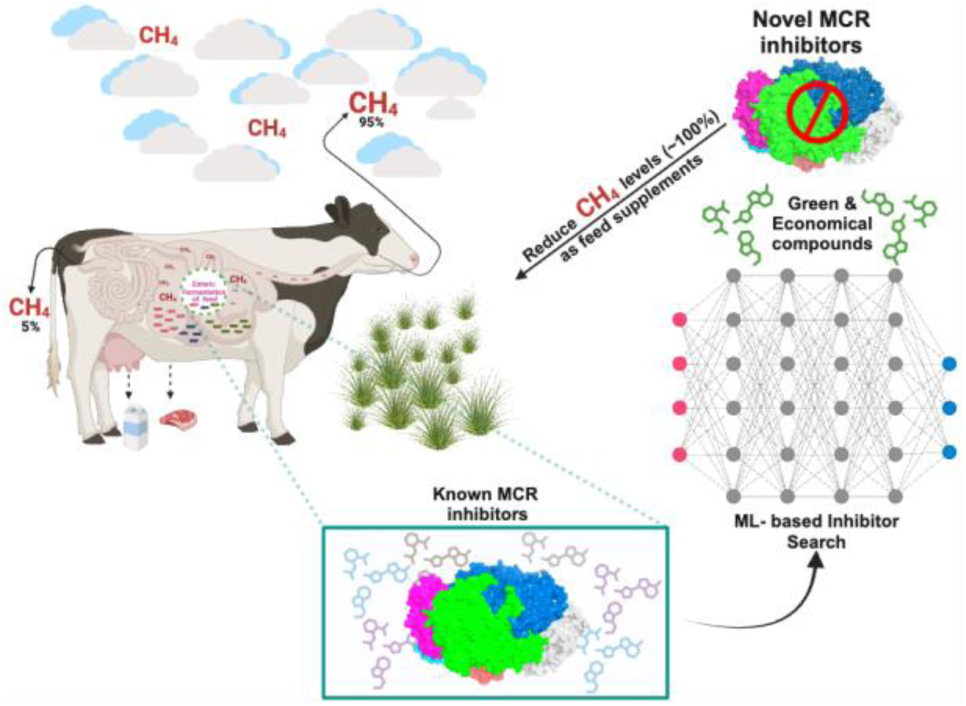

## INTRODUCTION

Greenhouse gases (GHGs) are atmospheric gases that possess the potential to absorb and retain infrared radiation in the atmosphere, hence trapping heat and causing a rise in temperature of the earth’s surface ^1,2^. Prominent GHGs encompass carbon dioxide (CO_2_), methane (CH_4_), nitrous oxide (N_2_O), as well as a selection of fluorinated gases^1–4^. GHGs have been one of the world’s major climate change drivers over generations since their emissions degrade the atmospheric layer. This results in global warming due to anthropogenic activities. Inclusive of these activities are enteric CH_4_ emissions from ruminant livestock, the release of CO_2_ from fossil fuel use, land use change, and landfills.

According to the sixth assessment report by the Intergovernmental Panel on Climate Change (IPCC)^5^, there were 59 Gt of CO_2_-equivalence (CO_2_-e) emitted globally in 2019. Emissions from Agriculture, Forestry and Other Land Use (AFOLU) represented 22% of these emissions. Enteric CH_4_ emissions accounted for 5.1% of total global GHGs, 23% of AFOLU, and 27.2% of total CH_4_ emissions **(Figure 1a)**. As CH_4_ has a short atmospheric lifespan (approximately 12 years), in periods where emission rates are reduced to a large enough degree, there will be less atmospheric CH_4_, resulting in lower warming. Accordingly, rapidly declining CH_4_ emissions can reduce temperature equivalent to the removal of atmospheric CO_2_. As such, reductions in CH_4_ emissions represent the most promising means to address climate change in the short term ^6^. This nuance of CH_4_ emissions in general and the relative contribution of enteric CH_4_ to total CH_4_ emissions make enteric CH_4_ mitigation particularly important ^7,8^.

**Figure 1.**
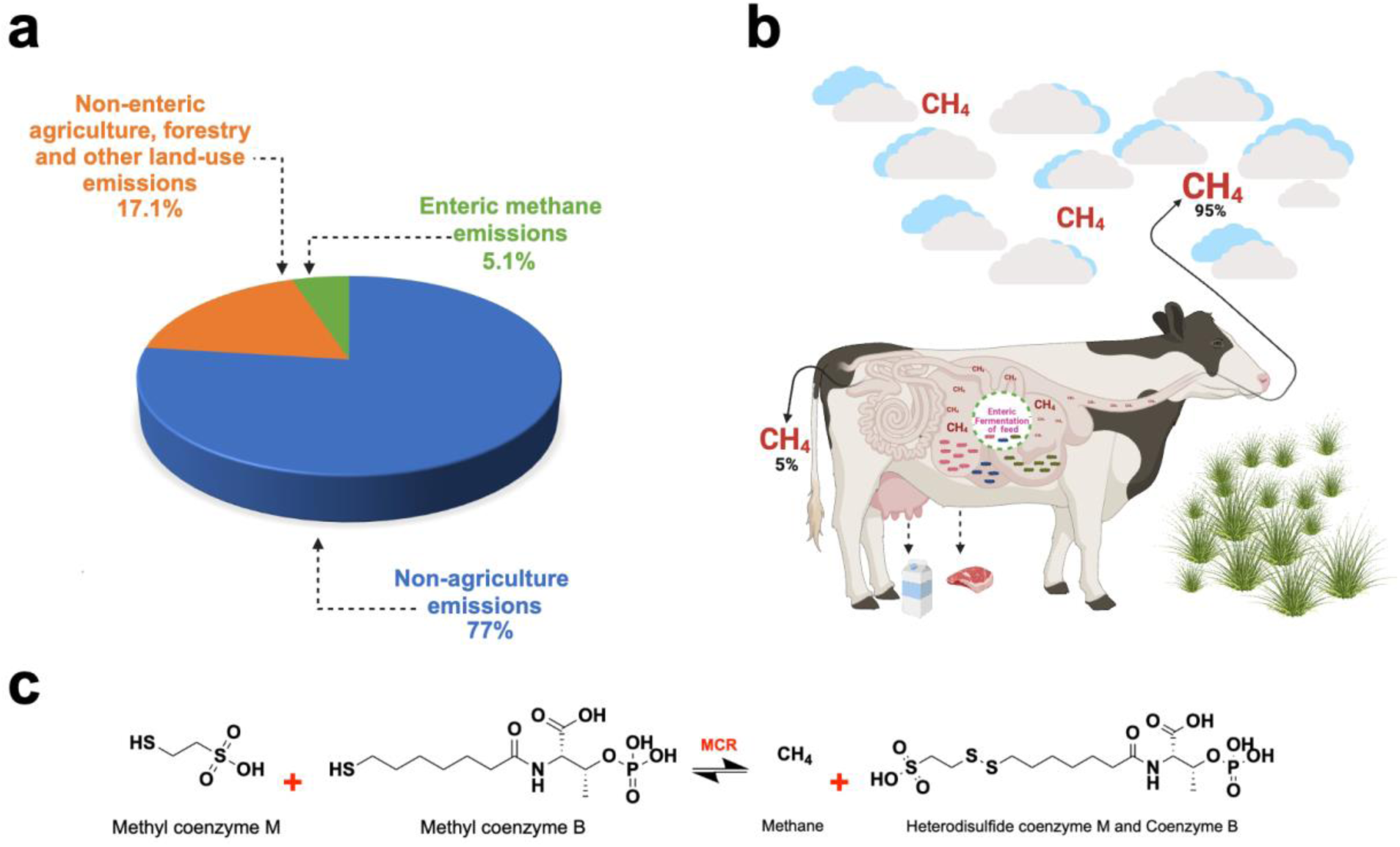
A comprehensive schematic illustrating the distribution of greenhouse gas emissions, focusing specifically on methane and detailing its biochemical synthesis and release mechanisms. **a)** Global representation of GHGs emissions with distributions centered on methane by sector as gathered from literature **b)** The entire enteric fermentation of carbohydrate (cellulose) feed as a mechanism of methane release. **c)** Biochemical reaction and the rate-limiting step in enteric methane synthesis catalyzed by MCR enzyme.

Due to increasing production efficiency, the carbon footprint of milk production was reduced by 40%, from 33.6 to 19.9 g CH_4_/kg milk in recent years, and reductions of 16.3% of enteric CH_4_ per unit of beef produced for 2007 relative to 1977 ^9^. While these reductions in carbon footprint from the dairy and beef industry are commendable, recent pledges of carbon neutrality by industries and companies have increased since the Paris Climate Agreement^10^. These types of commitments require reductions in absolute emissions rather than reductions in emissions per unit of product. Accordingly, enteric CH_4_ mitigation is highly needed by both the dairy and beef industries.

Methanogenesis is methane biosynthesis, irrespective of its emission source. Methane is a key natural secondary metabolite of enteric fermentation in the rumen of ruminants upon the digestion of consumed feed ^11^. Conditions favoring the production of enteric CH_4_ are designated to achieve homeostasis in the presence of excess hydrogen for maximum energy production. Methanogens (methanogenic archaea) are the predominant mediators of methanogenesis within the rumen. In agreement with the above, a study reported methanogens are influenced by other microbial members, primarily bacteria ^12^. Methanogenic interactions with bacteria, fungi, and protozoa influence enteric fermentation, the main metabolic reaction that leads to CH_4_ production. Therefore, methanogens represent a key target for investigating metabolic processes for CH_4_ mitigation.

Significantly, enteric CH_4_ production has been a conventional marker for farming productivity as CH_4_ is an associated product for carbohydrate utilization in ruminants. The quest for essential and volatile fatty acid production in livestock dietary metabolism has leveraged this gross implication of CH_4_ production in the four-chambered stomach of herbivorous grazing mammals^13^. As a natural result of excess hydrogen production in ruminants, CH_4_ is released into the atmosphere through either eructation (95%) or flatulence (5%)^1^ (**Figure 1b**). Following the stepwise biochemical reaction of CH_4_ biogenesis in ruminants, the enzyme Methyl Coenzyme M Reductase (MCR) produced from methanogenic archaea plays a key role. MCR catalyzes the final but rate-limiting step between methyl-coenzyme B (CoB-HS) and methyl-coenzyme M (CH_3_-S-COM) to release a heterodisulfide Coenzyme M and Coenzyme B (COM-S-S-COB) and CH_4_ as products^14^ (**Figure 1c**). The entire biochemical process is labeled methanogenesis for reference^15^.

Methanogenesis mitigation strategies and approaches have been conceptualized, designed, and deployed for a green and CH_4_-reduced ecosystem. Currently, several CH_4_ mitigation strategies are being explored by the agricultural sector. Options such as increasing feeding levels, decreasing dietary forage-to-concentrate ratios ^16,17^, and improving feed quality and digestibility have been promising options. However, these strategies often reduce enteric CH_4_ emission on a per product produced basis and have only demonstrated reductions by around 16.3% ^18^. Mitigation options that reduce absolute emissions have also been investigated and include genetic and breeding selection^19^, feeding tanniferous forages ^20^ providing electron sinks ^21^ and supplementing fat ^16,17^. These options have been shown to reduce enteric CH_4_ emissions by around 10% ^18^.

The mitigation options that have demonstrated the largest enteric CH_4_ mitigation potential are direct methanogenesis inhibitors. These include 3-nitrooxypropanol (3-NOP) and bromoform (CHBr_3_)-containing seaweeds *(Asparagopsis* spp.). 3-nitrooxypropanol has been shown to reduce enteric CH_4_ by 25-30% ^22^ and *Asparagopsis* seaweeds have reduced enteric CH_4_ by 80-98% ^23,24^. 3-nitrooxypropanol and CHBr_3_ from red seaweed have been suggested to inhibit methanogenesis by competitively binding and providing an agonistic effect on CH_3_-S-COM hence hindering the final and rate-limiting step in enteric methanogenesis ^1,25,26^. More specifically, halogenated compounds such as CHBr_3_ competitively displace other natural substrates that tend to interact with the Ni(I) ion of F_430_ coenzyme M. This results in methyl transfer inhibition and a reduction in CH_3_-S-COM mediated CH_4_ release^26^.

Amongst the methanogenesis inhibitors investigated and implemented, a data-driven deep-dive with precise molecular modeling of the atomic-level biochemistry of these inhibition mechanisms has remained largely elusive. Empirical approaches thus far have not provided enough biochemical information in order to design novel inhibitor molecules which can posit a high affinity of binding to rumen MCR bound to its cognate cofactor F_430_ aside 3-NOP^1,2^. Here, we employ *in silico* approaches to interpret the stoichiometric ratios (i.e., biophysical flooding) and binding affinities (i.e., biochemical trapping) of all well-documented inhibitor molecules against the redox-active nickel (Ni(I)) tetrahydrocorphin, coenzyme F_430_ cofactor of MCR.

## METHODS

### Selection of Methanogenic Protein Structure and Inhibitor Compounds

A data mining sweep through reported literature was performed encompassing Google Scholar, GenBank ^27^, and UniProt ^28^ databases to pinpoint the methanogenic archaea Methyl-coenzyme M reductase enzyme responsible for enteric CH_4_ biosynthesis. Studies focused on the biochemical mechanism of the MCR enzyme, inhibitor molecules, and structural insights of both MCR and the inhibitor molecules were shortlisted. Based on the structural insight, a high-resolution, X-ray diffracted crystallized MCR (PDB Accession ID: **5G0R**) was identified^2^ and downloaded from the Research Collaboratory for Structural Bioinformatics Protein Data Bank (RCSB PDB)^29^. Protein structure visualization, characterization, and determination of active site residues within a 5Å distance from the cofactor F_430_ were investigated using PyMOL^30,31^. A library of inhibitor molecules was downloaded from PubChem^32^ after a deep literature search for inhibitor molecules with or without experimental data from the above-mentioned literature databases. The MolView server^33^ was used to generate structures for inhibitors that were not available in PubChem.

### Molecular docking and molecular dynamics (MD) simulations

Molecular docking and MD simulations were conducted to explore further insights into the binding poses and proximities for CH_4_ inhibition amongst selected 16 individual inhibitor molecules with the Ni(I) of F_430_ cofactor of MCR. Rigid molecular docking was performed using AutoDock Vina^34^ to explore the binding interactions of the selected inhibitors out of a library of literature-derived small molecule compounds and the cofactor F_430_ of MCR (PDB ID: 5G0R). Protein and ligand preparation steps were conducted using AutoDockTools^34^. Using the gradient-based local search genetic algorithm built in AutoDock Vina ^34^, the docking energy scores and rankings of binding poses of each inhibitor molecule to the active-site of the MCR enzyme were obtained. Illustrations of inhibitor-MCR complex were generated using PyMOL^30,31^. Molecular dynamics of the respective top-scored conformations of MCR-cofactor F_430_-anti methanogen ternary complexes were set up using GROMACS 2023 macromolecular modeling package with CHARMM36 forcefield^35^ (see SI for details).

### Stoichiometric ratio and binding affinity analysis

The stoichiometric ratio and distribution of inhibitor molecules within the catalytic groove of MCR at the surface of cofactor F_430_ were analyzed. All ligands’ poses within an electron transfer range (<5Å) with bound Ni(I) of the cofactor F_430_ were selected. The number of such poses for each inhibitor represents the maximum biophysically permissible stoichiometric ratio against inhibitor molecule type.

### Structural comparison of MCR inhibitors with ruminant specific metabolite databases

The 16 inhibitors explored against MCR enzyme were compared for similarities in molecular fragments within two ruminant specific metabolite databases - a) Milk Composition Database (MCDB) ^36^ and b) Bovine Metabolome Database (BMDB) ^37^, containing 2,360 and 51,682 entries, respectively. The structural information of metabolites was downloaded in Structure-Data File (SDF) format and further processed to obtain canonical Simplified Molecular Input Line Entry System (SMILES) representation using RDKit^38^. These SMILES strings were used as input for a GNN to generate molecular embeddings, providing a standardized and machine-readable representation of the complex molecular structures present in milk and bovine metabolites. Initially, the RDKit cheminformatics package was utilized to extract each metabolite’s atomic identities and structural information into features as nodes and edges. These features were then passed into a simple GNN architecture containing 58 input neurons, corresponding to the different atoms present in the structure databases, a hidden layer with 64 neurons, and 128 output neurons. This GNN framework generated molecular embeddings as tensors with dimensions (N×128), where N represents the number of atoms in each metabolite, and 128 is the dimensionality of the embedding space. These tensors were subsequently averaged across the atomic dimension to produce a unified 128-dimensional vector representation for each molecule. The high-dimensional embeddings were reduced to two dimensions using t-distributed Stochastic Neighbor Embedding (t-SNE). t-SNE parameters were optimized, with perplexity set to the minimum value between 30 and the total number of molecules in the database. This dimensionality reduction facilitated the visualization of molecular relationships, enabling the identification of structural similarities and potential functional associations among the metabolites.

### Validation of clustered potential inhibitors via Tanimoto chemical similarity analysis and Haddock

We employed Tanimoto similarity analysis, utilizing Morgan fingerprints, to assess the similarity between selected metabolites and the set of MCR inhibitors^39,40^. Bovine metabolites were categorized into groups of two, those with the highest similarity (Likely Inhibitors Molecules; LIMs) and those with the lowest similarity (Unlikely Inhibitor Molecules; UIMs) with the 16 known MCR inhibitors. Categorization was done based on clustering proximity. Additionally, we performed molecular docking studies using HADDOCK on five of the nearest and five of the farthest metabolites, targeting the enzyme MCMI reductase^41^. The active residues from the enzyme were selected for docking with the chosen metabolites (**as detailed in S1 Figure**). We predicted the expected membrane permeability of randomly chosen 16 Likely and Unlikely Inhibitor Molecules (LIMs/ UIMs), using an established supervised machine learning protocol^42^. Herein the SMILES representation of each molecule is one-hot encoded using an encoder-decoder setup and mapped to the respective membrane permeabilities (preferentially trained on colorectal adenocarcinoma cell membrane; Caco-2). The above protocol is housed within the KNIME suite of ML platforms^43^.

## RESULTS AND DISCUSSION

### MCR from Methanothermobacter marburgensis and diverse inhibitors identified as key targets for methanogenesis inhibition

We selected an x-ray defined crystal structure of MCR protein with a Ni-methyl species that is a proposed catalytic intermediate in MCR. The methyl group of methyl-coenzyme M stated usually situates at least a 2.1 Å proximal to the Ni(I) of the MCR coenzyme F_430_ for a successful catalysis to materialize. A rearrangement of the substrate channel has been posited to bring together substrate species; however, Ni (III)-methyl formation alone does not lead to any observable structural changes in the channel ^2^. Given this, studies with biochemical and structural analysis of the MCR from *Methanothermobacter marburgensis* were focused upon with the assumption that the last step of CH_4_ production in ruminants is the rate-limiting step of methanogenesis ^44^. A recent experimental study ^2^ of the inhibitory properties of 3-NOP with the 3D structure of MCR (PDB ID: 5G0R) deciphered at a high resolution of 1.25 Å was selected for our study. In agreement with previous literature ^44–46^, the MCR protein selected is a 273 kDa hexameric protein (**Figure 2**) with two catalytic subunits that are 50Å apart. The MCR protein has a deep active site pocket with a substrate groove that runs ∼ 30Å from to the protein’s surface^1^. The activity of MCR, as reported by computational analysis from experimental data^44,47^, demonstrated the enzyme remains active only when its Ni ion in the tetrapyrrole derivative of the cofactor F_430_ has a +1-oxidation state, therefore catalyzing the last CH_4_-production step of methanogenesis in the rumen of livestock such as cattle, sheep, and goats^1,48,49^.

**Figure 2.**
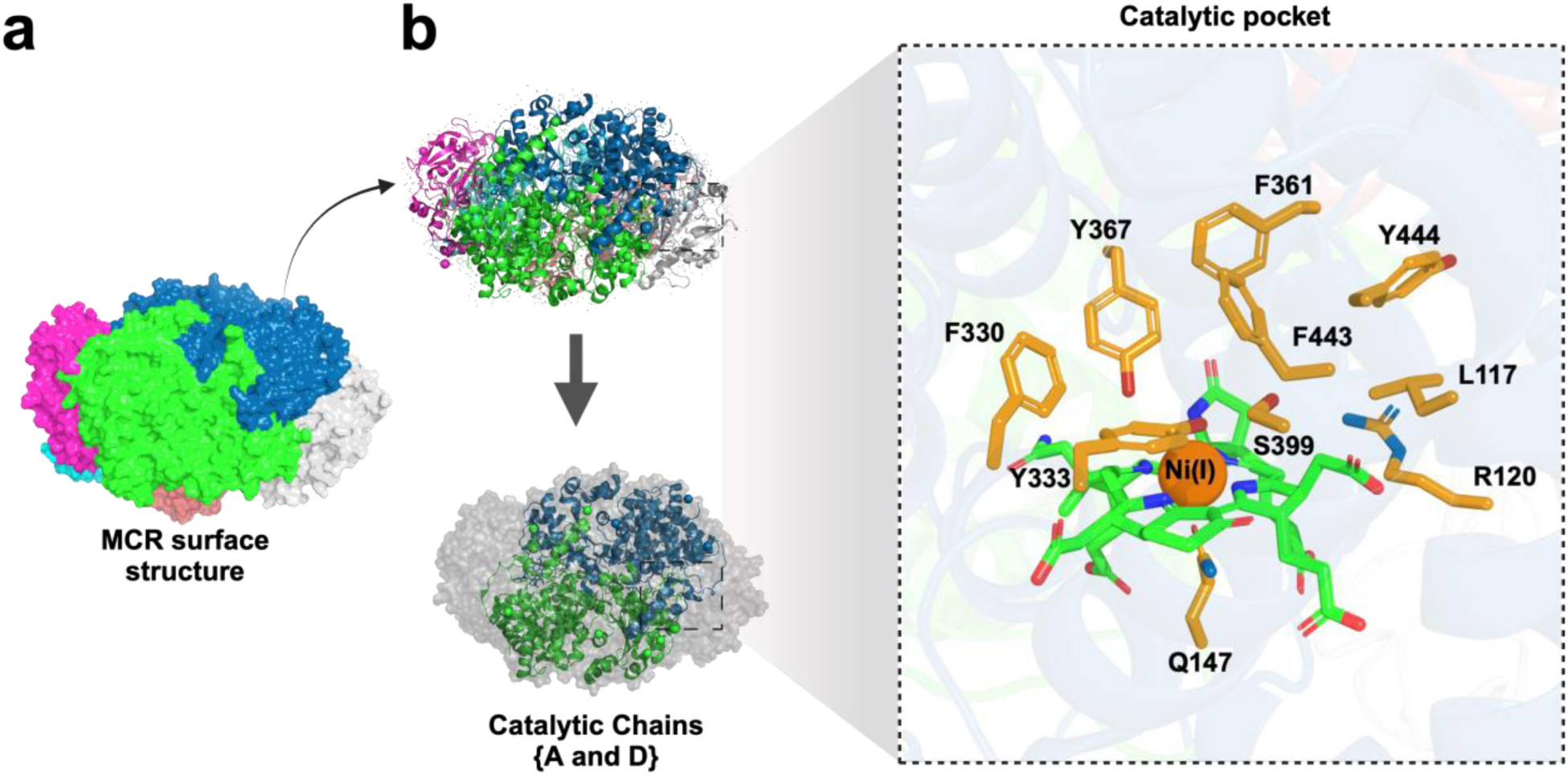
Illustration of the crystal structure of Methyl Coenzyme M Reductase (MCR) (PBD accession ID: 5G0R) from *Methanothermobacter marburgensis* and the six-chain hexameric complex. **a)** Each chain of MCR crystal structure has been indicated with six colors. **b)** Catalytically active chains (A and D) of MCR are shown in green and blue, while other non-catalytic chains are shown in gray. The location of the cofactor F430 in the enzyme structure for both catalytic chains are indicated.

All literature-based reported inhibitor compounds for enteric methanogenesis inhibition were collected. Sixteen distinct molecular compounds were selected, including statins, pterins, nitro-ol/esters, Coenzyme-B analogs (COBs), and CHBr_3_ (see **Figure 3**). Three statins (atorvastatin, rosuvastatin, and simvastatin), four nitro-ol/esters (2-nitroethanol, 2-nitropropanol, 3-nitropropionate and 3-NOP), five Coenzyme B analogs (COBs) (N-5-mercaptopentanoylthreonine phosphate: CoB5, N-6-mercaptohexanoylthreonine phosphate: CoB6, N-7-mercaptoheptanoylthreonine phosphate: COB7, N-8-mercaptooctanoylthreonine phosphate: CoB8, and N-9-mercaptononanoylthreonine phosphate: CoB9, and three Pterins (pterin B53 (2,6-diamino-5-nitrosopyrimidin-4(3H)-one), pterin B54 (4-{3-[(2-amino-5-nitroso-6-oxo-1,6-dihydropyrimidin-4-yl)amino]propoxy}benzoic acid) and pterin B55 (2-amino-8-sulfanyl-1,9-dihydro-6H-purin-6-one)) were studied in comparison with CHBr_3_ using detailed molecular modeling and thermodynamic assessment of binding interactions with MCR in the presence of cofactor F_430_.

**Figure 3.**
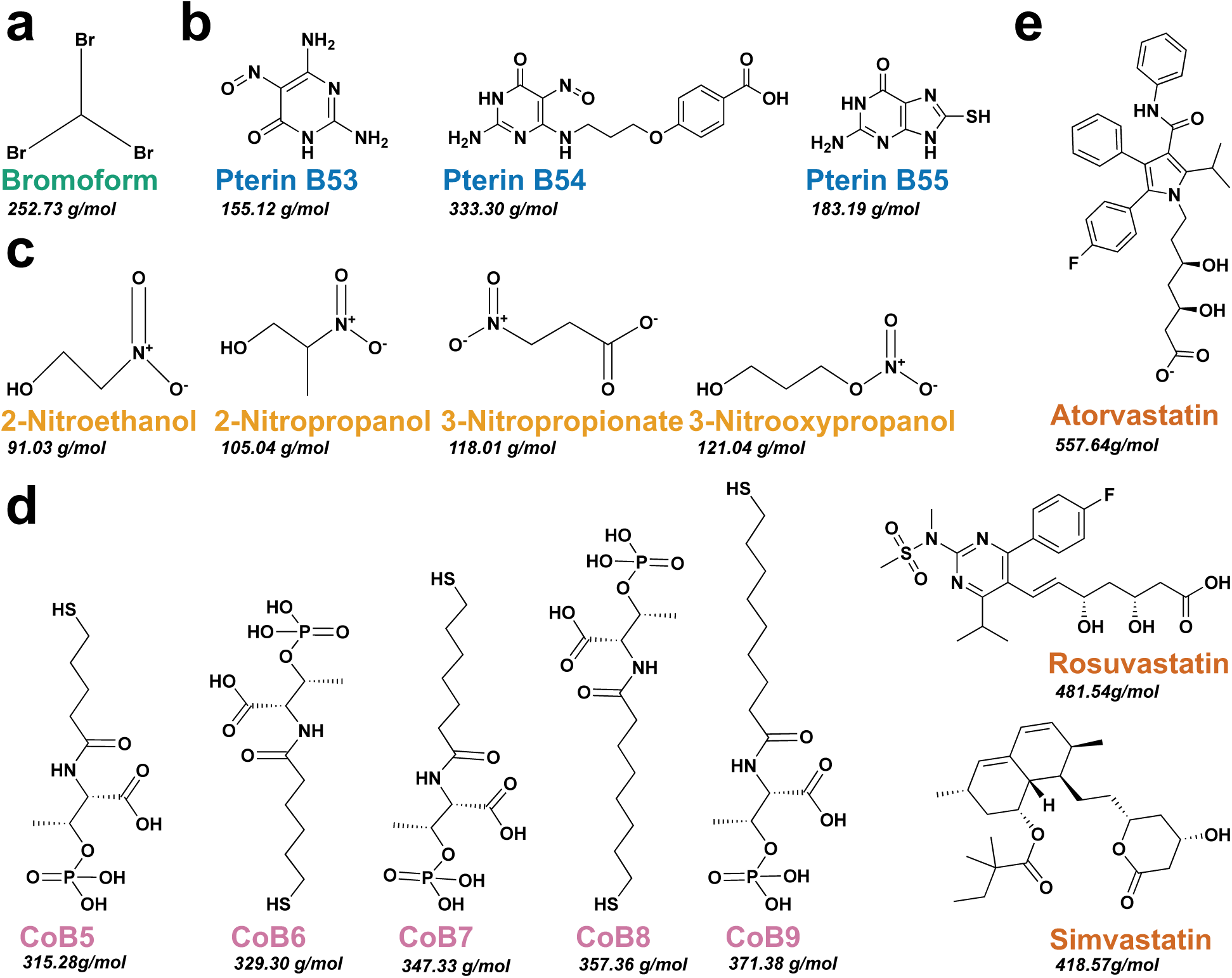
Representation of all selected anti-methanogenic molecules (inhibitors) structures adopted for this study. **a.** Bromoform molecule. **b.** Group of Pterins. **c.** Group of Nitro-alcohols and esters. **d.** Group of Coenzyme B analogs. **e.** Group of Statins or HMG-CoA reductase inhibitors.

### Nitro-ol/ester compounds outperform other inhibitors in MCR binding affinity and stoichiometry

The top-scoring (strongest) binding poses were analyzed to evaluate the ligands’ binding affinities, interactions, and potential binding modes with no superimpositions within 5Å. No superimposition criterion was imposed to infer the maximum number of inhibitor molecules that can simultaneously invade and yet remain biochemically bound within catalytic distances of cofactor F_430_ within the MCR enzyme pocket. The number of inhibitor molecules thus obtained is a representation of the maximum permissible stoichiometry of the inhibitor on a per-molecule basis with the MCR enzyme. Consequently, the inhibitor molecule poses that were accounted for were the ones within the electron transfer range with the Ni(I) of the tetrapyrrole of F_430_ in MCR. The least likely inhibitor molecule from the binding affinities records were HMG-CoA reductase inhibitors (statins) with rosuvastatin, simvastatin, and atorvastatin having positive (i.e., overall repulsive binding interactions with MCR). They had +55.5, +54.7 and 79.7 kcal/mol binding energy scores, respectively – reflecting they are unlikely to stay bound and/ or inhibit catalysis sustainably, even though they are shape compatible for the MCR pocket and might temporally occlude the pocket. It is noteworthy that the MCR-binding affinity values observed with the statins numerically correlate (R^2^ = 0.82) with the molecular weights of each statin due to the tube-like shape of the binding pocket of MCR. Next, the coenzyme B analogs ranked as poor, albeit stable inhibitors from the affinity values from the top three docking poses per inhibitor, with CoB5 having the lowest affinity value (3.9 kcal/mol). However, the third best group of inhibitors was the pterins, with pterinB55 being the best amongst them at 2.17 kcal/mol, while the worst of that group was pterinB54 with an affinity binding of 15.52 kcal/mol. Best as desired, inhibitor molecules surfaced as the nitro-ol/ester group of molecules with mean affinity values ranging from −2.87 to −5.37 kcal/mol (see **Figure 4**). The CHBr_3_ molecule scored an average affinity value of 1.33 kcal/mol, with the best individual CHBr_3_ molecule having a 0.2 kcal/mol but stoichiometrically having three poses with no superimposition (**Figure 4**).

**Figure 4:**
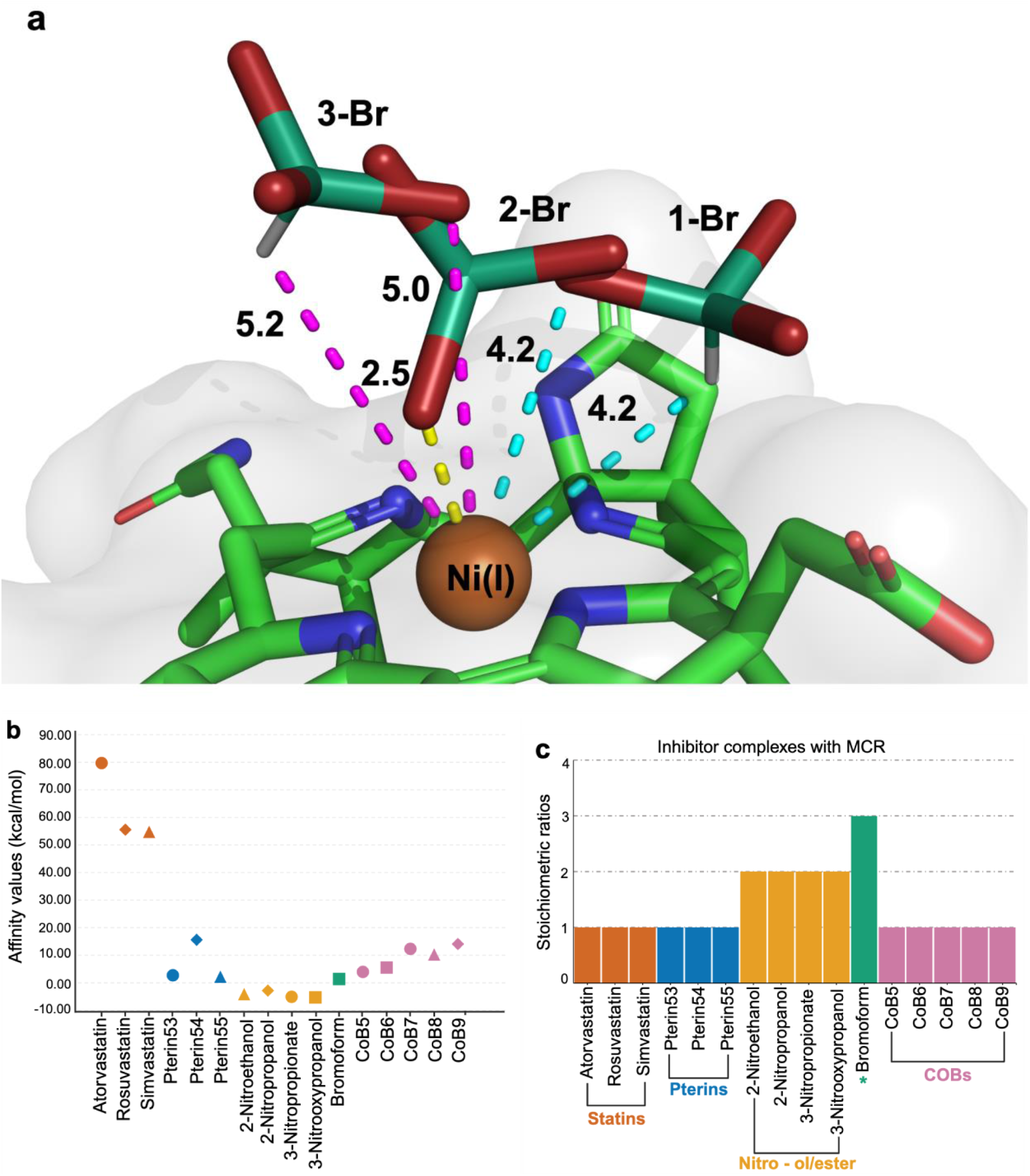
Illustration of all three selected poses of bromoform interacting with Ni(I) of F_430_ in MCR protein and graphical representation of the stoichiometric ratio of individual inhibitors docked to the active site of MCR enzyme in the close vicinity of F430. The dashed lines indicate the distances, in Å, between Ni(I) and bromoform. **Cyan**: for the distances of the first bromoform molecule. **Gold**: for the distances of the second bromoform molecule. **Magenta**: for the distances of the third bromoform molecule. The distances of other inhibitor molecules from Ni(I) are represented in supplementary information (see Figure S2-S17). **b.** Scatter plot representation of the mean binding affinity values of top three conformations of inhibitor molecules docked to F_430_ of MCR. **c.** Representation of all positive conformations of inhibitor molecules accurately posed within a 5Å range.

Energetics for each inhibitor (*in silico* affinity value scores) were calculated based on the best conformations docked at the active site. The relatively small CHBr_3_ and nitro-ol/ester compounds were observed to be comparable with each other and better than the other anti-methanogenic compounds due to their larger molecular size; however, fragments that interacted with F_430_ need to be analyzed further for more insights. Affinity values of each inhibitor (selected poses) were plotted for the compounds which are correlated with the stoichiometric ratio plot (**Figure 4**). Apart from CHBr_3_, which has more experimental evidence, nitro-ol/ester compounds could be stable enough for competitive inhibition. Molecular dynamics for these compounds warrant further investigation into their anti-methanogenic capabilities.

### Ni(I) ion mobility and steric clashes hinder stable MCR-cofactor F430 complex simulations

The MCR enzyme is a hexameric enzyme with two catalytic grooves, each guarded by three chains (**Figure 2**). The active form of cofactor F_430_ has a tetrapyrrole ring with Ni(I) held at its center. To reduce the computational cost without compromising the quality of MD simulation, we focused on one catalytic groove, which encompasses chains A, C, and D along with cofactor F_430_. The force field parameters for MCR are taken from CHARMM36, while cofactor F_430_ is parametrized using ATB ^50^. Equilibration of the three chains of enzyme along with the cofactor F_430_ with Ni(I) in an orthohedral TIP3P water box resulted in cofactor F_430_ moving out of the solvation box and Ni(I) moving away from cofactor F_430_ into the bulk solvent (**SI Figure A**). Selection of orthohedral simulation box is to reduce the solvent molecules with the aim to reduce computational cost. The undesirable shifting of cofactor F_430_ in orthohedral box may be a result of the edge effect due to poor solvation, hence we controlled it by using a cubic water box, with 2.8 times increase in number of solvent molecules. Nevertheless, the tendency of Ni(I) to behave as a solvent ion continued to pose difficulty in modeling MCR-cofactor F_430_ complex (**SI Figure B**). We attempted to control the relative movement of Ni(I) with respect to cofactor F_430_ by imposing movement restrictions, which resulted in unfeasibly unstable energy due to steric clashes. Adding inhibitor molecules to a non-equilibrated enzyme-cofactor complex further worsened the instability of the whole simulation system, resulting in an unphysical simulation box.

As atomic scale MD simulation of MCR enzyme-cofactor F_430_-inhibitor ternary complex is a challenging venture involving multiple steps of optimization, equilibration, and analyses involving a huge computational cost, we intend to implement the knowledge we gained in optimizing the simulation box for a future follow-up study to compare the structural and thermodynamic underpinnings of MCR inhibition ^51^.

### MCR inhibitors cluster together when compared with ruminant specific metabolite databases

Spatially adjacent molecules to the inhibitor cluster in the reduced-dimensionality space emerge as putative inhibitors or precursors of anti-methanogenic compounds. Since it is unclear what characteristics define a good inhibitor, as all 16 molecules are very different from each other in shape and chemistry, binding energy calculations can tell if a molecule is a good inhibitor, but this information alone is insufficient to design a new inhibitor. This necessitates the identification of common structural and chemical features that unify these 16 molecules while simultaneously distinguishing them when put in context with other bovine metabolites. Since the number of molecular features required to identify such a cluster is unknown due to paucity of data in experimental literature, we chose to use a latent encoder of molecular signatures using a graph neural network (GNN) whose encodings when projected onto a 2D space, exhibits clustering of these 16 molecules close to each other and disparate from others. While other functional clusters have not been investigated in context with bovine metabolism and signal transduction, we are able to ascribe the clustering of all these 16 validated anti-methanogenic molecules to represent the loci in the 2D t-SNE space as responsible for anti-methanogenicity (**Figure 5**).

**Figure 5.**
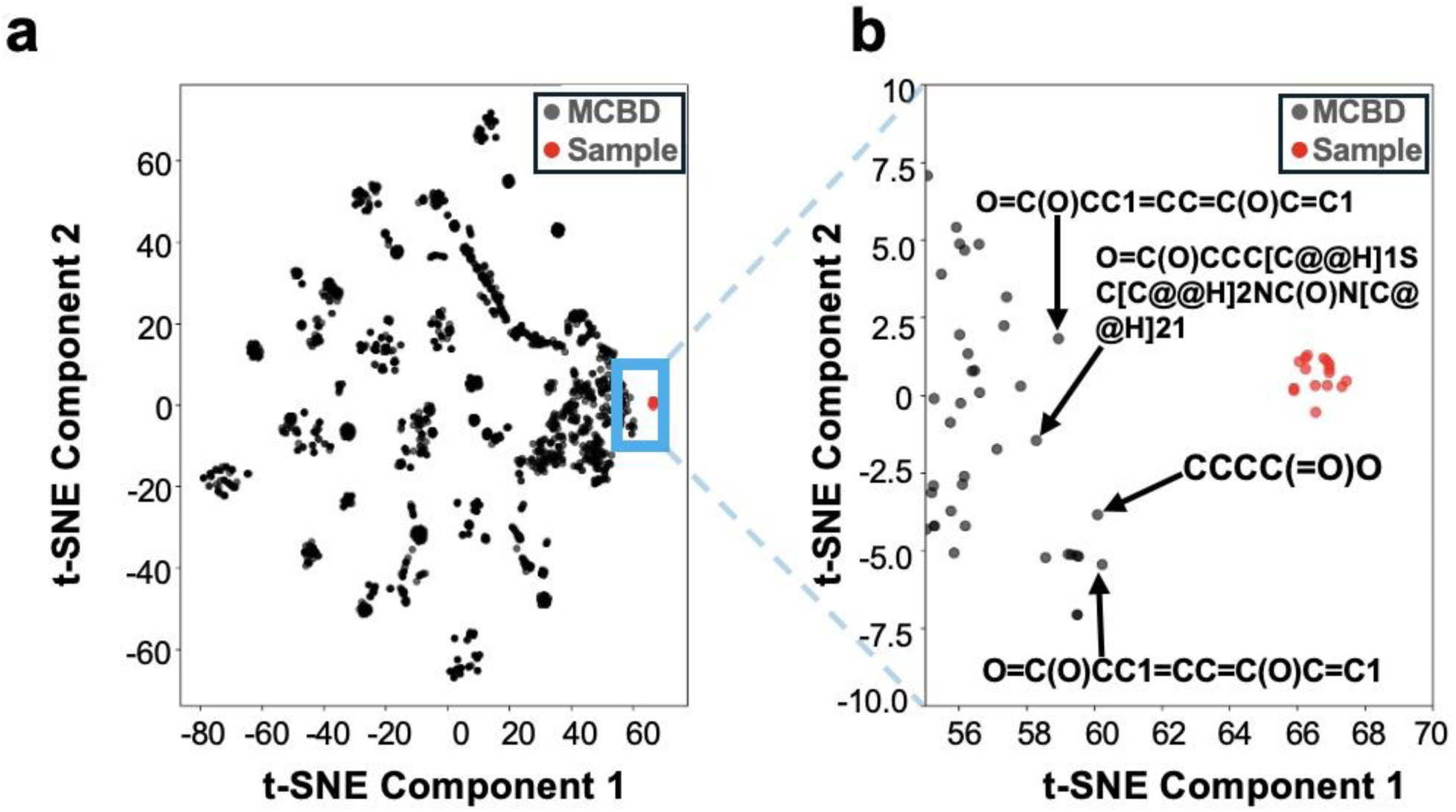
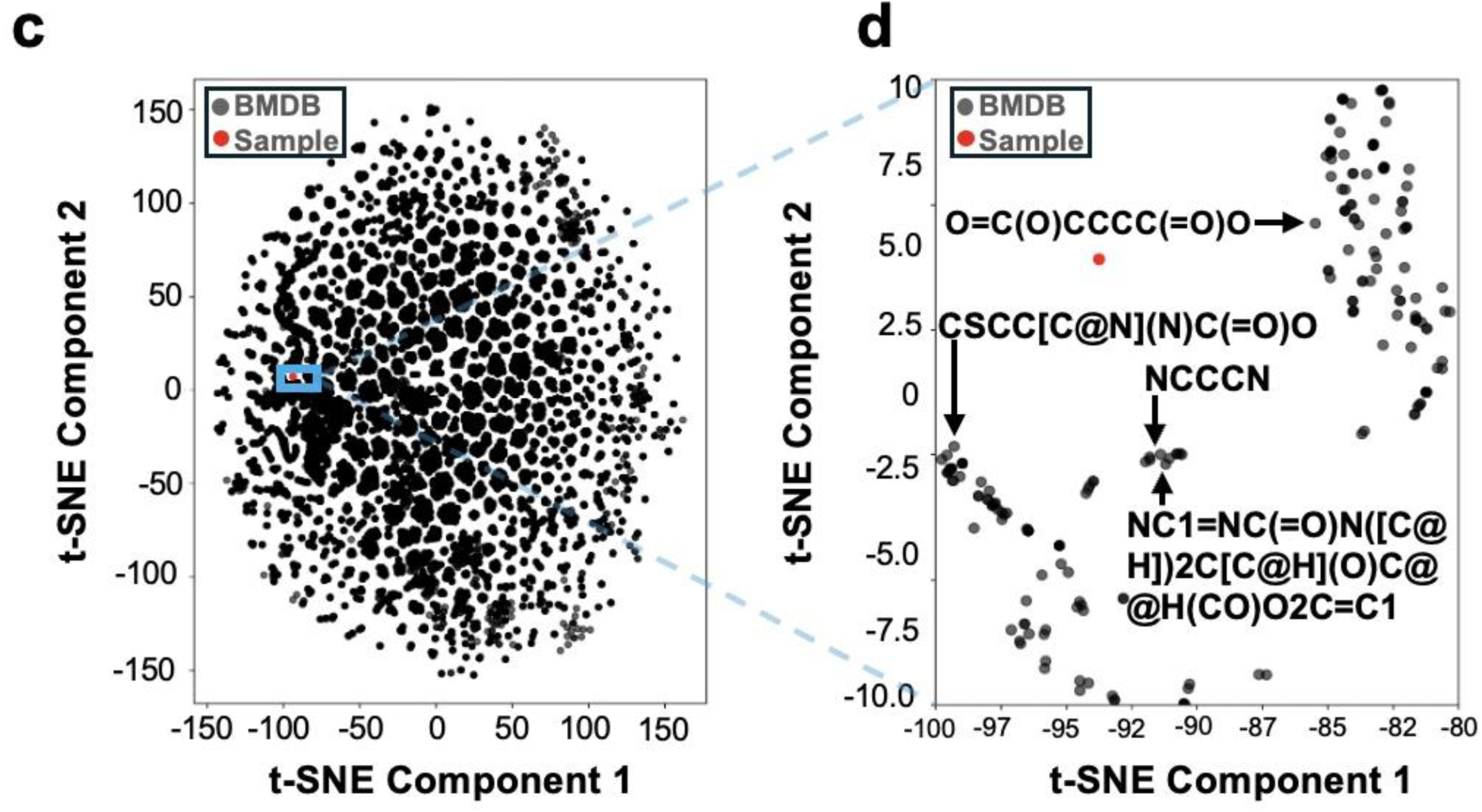
Two-dimensional t-SNE projection of molecular signatures reveals clustering of methanogenesis inhibitors. a) and b) Visualization of 16 known MCR inhibitors (Red) in relation to their four nearest neighbors (Black) selected from the Milk Composition Database (MCDB). c) and d) Similar visualization with four proximal metabolites (Black) identified in the Bovine Metabolome Database (BMDB).

Proximal molecules to this functional cluster from the two databases emerge as putative inhibitors or precursors to anti-methanogenic molecules. Notably, molecules such as butyrate, 2-hydroxybutyric acid, and biotin were identified as potential candidates. Previous studies in the field address the success of computational tools for the prediction of inhibitors for various enzymes. From the discovery of novel QoI fungicides for cytochrome b inhibition in *Peronophythora litchi* ^52^ and the successful elucidation of antimicrobials for downy mildew pathogenicity in cucumber using in silico docking ^53^. Over the period of advancement, the use of ML-based tools^54,55^ dominates the race of drug or ligand prediction after several successes. On this note, our team’s next steps are to leverage generative AI frameworks like Drug-large language models (LLM) or Chemistry42 in subsequent studies ^51^ to computationally predict potential inhibitors using the putative inhibitors as templates and couple it with *in vitro* inhibitor assays to test the efficiency of such predictions.

### Validation of clustered potential inhibitors via Tanimoto chemical similarity analysis and HADDOCK

We demonstrate that the LIMs (likely inhibitors) exhibited significantly higher Tanimoto similarity scores with the known sixteen inhibitor molecules compared to the UIMs (unlikely inhibitors) metabolites (**Figure 6 (a)** and **(b)**). We conducted a t-test that yielded a *p value* of 0.0003 indicating that LIMs have a significantly higher chemical similarity (Tanimoto score) to the known inhibitors, compared to UIMs. This provides interpretability to our neural clustering (**Figure 5**). The chemical similarity trends, however, did not correlate with the HADDOCK computational docking scores (i.e., binding free enthalpies) with the MCR enzyme, as distal metabolites (UIMs) often resulted in tighter MCR binding (**SI Table 3**). This can be ascribed to the lack of appropriate biochemical microenvironment in a static docking simulation which ignores entropic effects of solvent molecules (see details on attempted MD simulations; **Supplementary Information Figure S18**). The complexity of the dynamics of this quaternary system (an enzyme, a F430 cofactor, a Ni(I) metal ion, and an inhibitor) when interfaced with explicit water molecules becomes intractable as seen in our attempt to perform the MD simulation (due to the paucity of all appropriate non-bonded parameters). This even more alludes to the lack of fidelity in available docking protocols which are not poised to handle co-docking setups with more than two moving pieces. Despite the accurate identification of the key (active) residues involved (**S1 Figure**) in substrate stabilization, HADDOCK results were thus not contributive to explaining the true energetics of the system. Overall, these findings suggest that chemical similarity, as measured by Tanimoto scores, is likely to be a more reliable predictor of MCR inhibition potential than inhibitor binding affinity.

**Figure 6.**
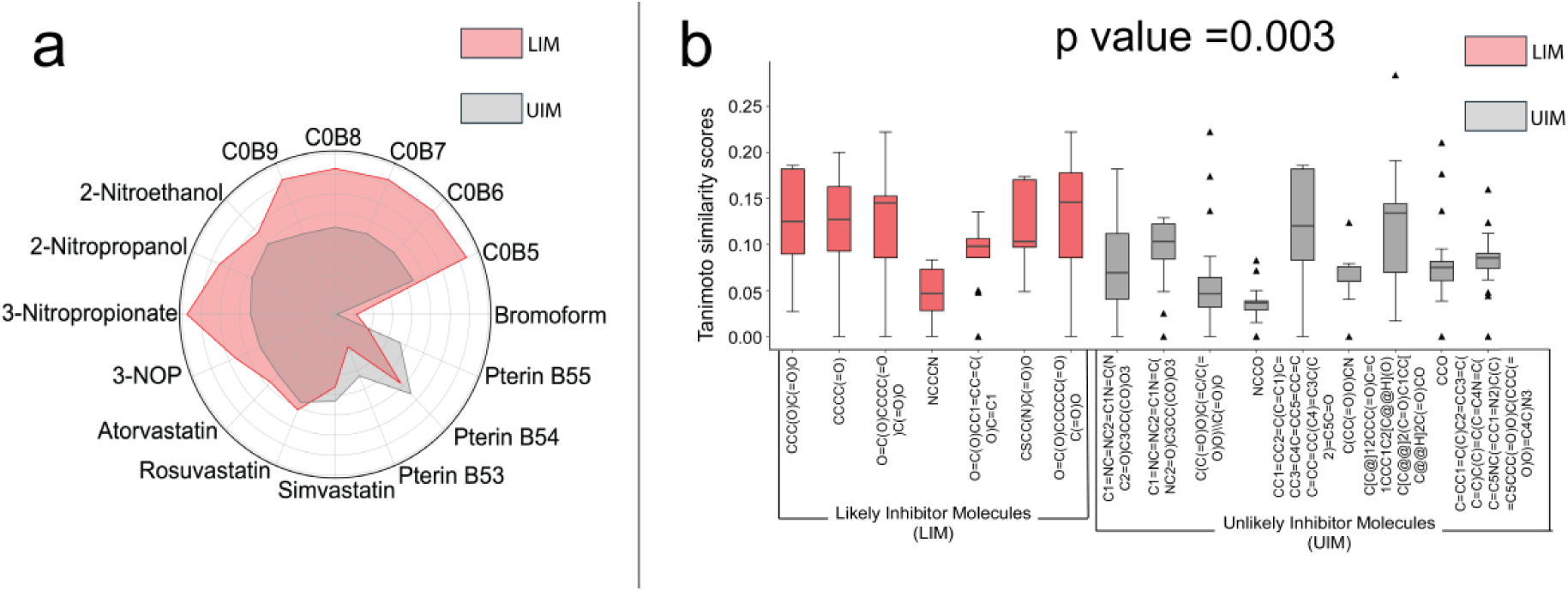
Tanimoto chemical similarity analysis between the LIM and UIMs relative to sixteen MCR inhibitors. (a) The sixteen inhibitors are represented at the periphery of the spider plot. The red-shaded area indicating the similarity of the proximal LIMs while the gray-shaded area represents the farther UIMs. b) Box plots illustrate the similarity of seven LIMs and nine UIMs relative to the 16 known MCR inhibitors. The red boxes represent LIMs, while the gray boxes represent UIMs. A *p value* of 0.003 indicates that the LIMs exhibit a statistically significant higher similarity to the known sixteen inhibitors compared to UIMs.

### Membrane permeable metabolites are likely to inhibit the methane emission in ruminant

MCR is mostly associated with the membrane^56^, and recent findings confirm its localization near the cytoplasmic membrane^57^. This indicates the necessity of membrane permeability for effective inhibition. For instance, bromoform and 3-NOP are established MCR inhibitors^26,58^. Bromoform is known to penetrate cell membranes rapidly, achieving diffusion within nanoseconds at low concentrations^59^. 3-NOP has been shown to significantly reduce methane emissions in dairy cows, leading to its approval for commercial use by the FDA^58,60^.

In our analysis, bromoform was predicted to exhibit high membrane permeability, albeit with low computational prediction confidence. Among putative inhibitors (without further property screening) candidates like CSCC(N)C(=O)O (S-methyl cysteine) and CCC(O)C(=O)O (4-hydroxybutyric acid) emerge as highly permeable with high confidence (**Table 1**). They have high chemical similarities (median Tanimoto scores ∼0.10, ∼0.13 respectively) with the known sixteen metabolites. While S-methyl cysteine is a known anti-oxidant, anti-inflammatory^61^, and is biologically regarded as safe^62–64^ and hence a promising target for experimental testing, 4-hydroxybutyric acid (Drugbank id: DB01440) is known to be a therapeutic drug and can lead to cytotoxicity^65^ above when administered beyond threshold. This makes the latter a less promising experimental target. It indicates the necessity to build additional bio-aware filters into computational predictive models beyond chemical similarity, membrane permeability and ability to approach Ni(I) before taking computationally predicted molecules to experimental testing for MCR. This is exemplified, as 3-NOP is predicted to have low permeability (even though with low confidence) (**Table 1**) which is contrary to experimental knowledge. Given its established use as a commercial feed additive, 3-NOP should have exhibited high membrane permeability in our predictions. One potential reason could be lack of 3-NOP-type molecules in the existing databases, making the prediction low confidence anyway (**Table 1**). Therefore, there is a clear need for a more precise Caco-2 membrane permeability predictor with biochemical awareness. Future work may involve developing advanced models, such as nonlinear regression or gradient-boosted trees^66^, leveraging data on 511 known metabolites with permeabilities across 11 representative membranes.

**Table 1:**
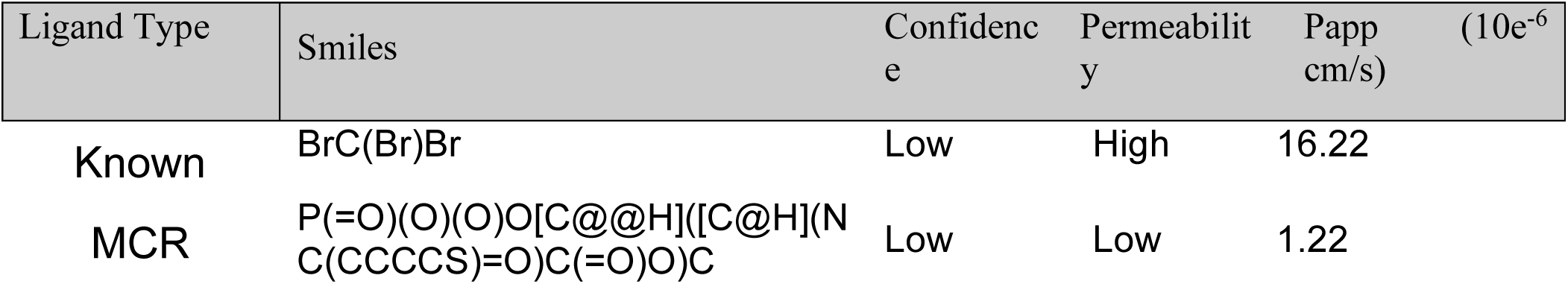
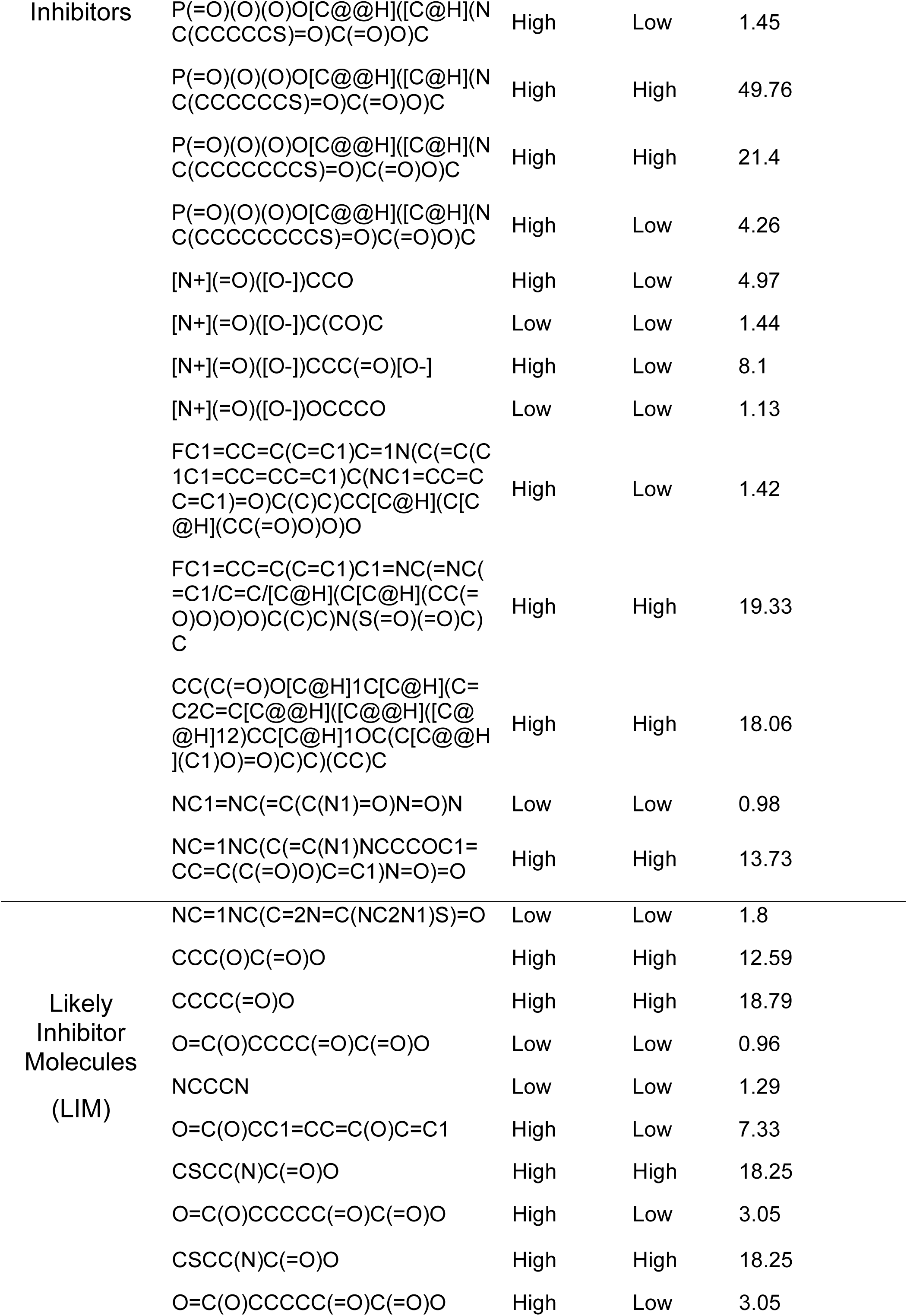
Predicted Membrane Permeability and Confidence Levels of MCR Inhibitors and Near Metabolites Based on SMILES Codes.

## CONCLUSION

MCR enzyme inhibition is considered a direct strategy to reduce CH_4_ emission from ruminant livestock. Here, we computationally compared 16 small molecules reported to be explored as MCR inhibitors. Through molecular docking, we showed that CHBr_3_ and nitro-ol/ester compounds have a higher affinity to bind to cofactor F_430_ in the active site of MCR compared to statins, pterins, and COBs. In this study, we revealed that the reaction dynamics and the overall mechanistic understanding of the inhibition process is greatly influenced by the stoichiometry of the inhibitors in the active site. Specifically, the presence of three bromine atoms in bromoform makes it a highly effective halogenated compound for competitively inhibiting the interaction of natural substrates with the Ni(I) ion in the F_430_ cofactor in MCR enzyme. Notably, inhibitor stoichiometry does not only dictate the binding affinity as a factor for methane inhibition but also the extent of methyl transfer inhibition and, consequently, the reduction in methane (CH₄) release. In this study, we demonstrate that the stoichiometry of the inhibitors in the active site, as deduced from the non-superimposing docking poses within the active site groove, is directly proportional to the size of the inhibitor. It can be interpreted that smaller inhibitors have higher flooding effects within the active site. The GNN-powered t-SNE clustering indicated that all the 16 inhibitor molecules explored in this study have inherent similarities among themselves when compared to ruminant specific metabolites and reveal some potential candidates from these databases as anti-methanogenic agents and their precursors. Lastly, the challenges in setting up an atomic scale MD simulation box with MCR enzyme-cofactor F_430_ with an electrostatically bound Ni(I)-inhibitor ternary complex is discussed, indicating the importance of optimizing each component of the ternary complex solvated in a solvent box big enough to ultimately house all the components.

## ASSOCIATED CONTENT

## Supporting Information

The Supporting information is compiled and available free of charge at the link to be added later.

## AUTHOR CONTRIBUTIONS

The project was conceived by RC. The simulations and analyses were set up and performed by RA. KAS and RA performed the molecular dynamics simulations while machine learning model training and prediction of prospective inhibitor molecules were done by MSN and AB. SN helped in data collection. SD conducted Tanimoto similarity analysis and haddock. TJM provided a valuable discussion on the competitive inhibition of enzymes, which guided the study. MB, NF, and JK provided valuable feedback which helped in designing the study. RA and RC wrote the manuscript. All authors helped in editing the manuscript. No authors declare any competing interests. All authors agree with this final version of the manuscript.

## ACKNOWLEDGMENT

R.C. acknowledges support through Iowa State University Startup Grant, Building A World of Difference Faculty fellowship, CIRAS Applied Mini-Grant, and NSF 22-599, EPSCoR RII Track-1, Award Number DQDBM7FGJPC5 for partially funding this study. R.A. was funded by NIH R35GM143074 to T.J.M. T.J.M. is also supported by the Karen and Denny Vaughn Faculty Fellowship. The authors thank Bibek Acharya for helping set up the force fields for the inhibitors to be compatible for molecular dynamics simulations. We also acknowledge the use of Google Scholar for the significant literature search process as it greatly contributed to collation of relevant references used in this study.

## ABBREVIATIONS

CCR2: CC chemokine receptor 2
CCL2: CC chemokine ligand 2
CCR5: CC chemokine receptor 5
TLC: thin layer chromatography

