## Supplemental File 1 for "Exploring putative enteric methanogenesis inhibitors using molecular simulations and a graph neural network"

**Active site description of methyl coenzyme m reductase (MCR)**

MCR enzyme catalyzes the reduction of methyl coenzyme M (CoM-SH) in the gut of rumen to produce methane ($CH_4$) as a natural means of reducing excess hydrogens for energy production. This reaction only proceeds in the presence of a key oxidized state of Ni, a component of cofactor $F_{430}$ situated in the active site groove of MCR enzyme. The MCR enzyme has a hexameric chained crystal structure as shown earlier with two symmertric catalytic sites. One active site situated in chain A while the other in chain D. As such, the catalytic site studied in this study was based on chain A yet the groove which is made up of chain A, D and C were jointly studied.

For visualization of key active site residues that aid in catalysis or form the active pocked were determined by visualization of the MCR enzyme (PDB ID: **5G0R**) in PyMOL software. As shown below in figure S1, the active site residues with both lateral and top views as surfaces.

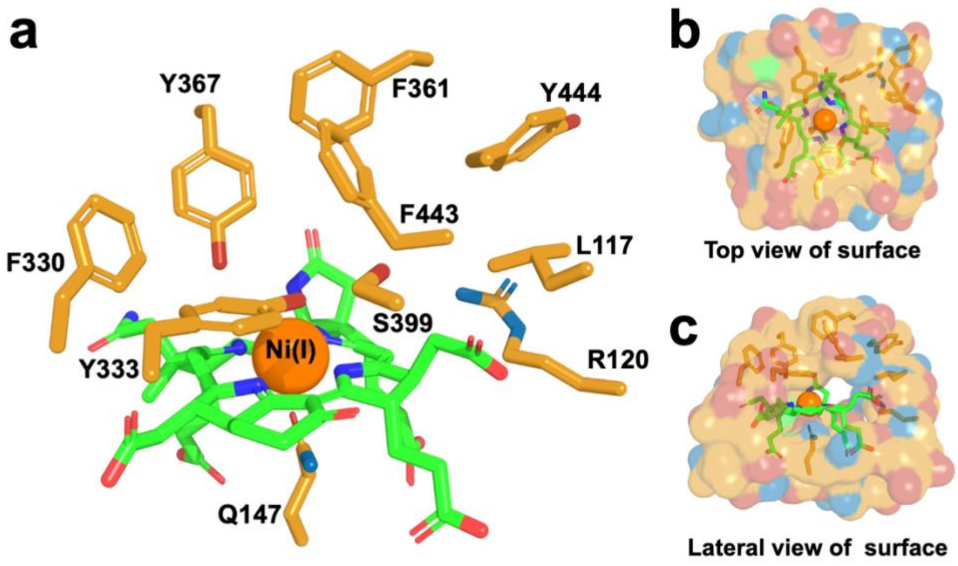


**Figure S1**: Representation of key active site residues of methyl coenzyme m reductase (MCR) enzyme in 5 Å proximity with Ni(I) of cofactor F430 along with surface views.

### Molecular Docking

Selected sixteen inhibitor molecules deduced from literature as reported were docked using AutoDock Vina and AutoDock Tools jointly. As following the rigid docking procedure, MCR enzyme was kept rigid while inhibitor molecules were made to freely move in the system. A grid box of 128x128x128 was used with focus on the cofactor $F_{430}$ of MCR. Binding poses from the results were screened with a non-superimposing criterion as well as poses within 5 Å distance from the Ni(I) of $F_{430.}$ Below are the top selected poses of inhibitor molecules within 5 Å proximal range of Ni(I) of cofactor F30 within the active site groove of MCR used in this study with their affinity values (**see table S1.0**).

**Table 1: Energy scores of top 3 binding poses and their mean for all inhibitors.**

| Type of Molecule | Energy scores | | | |
|---|---|---|---|---|
| | Pose A | Pose B | Pose C | Mean |
| Atorvastatin | 79.7 | 79.7 | 79.7 | 79.7 |
| Rosuvastatin | 55.3 | 55.5 | 55.7 | 50.5 |
| Simvastatin | 53.6 | 54 | 56.5 | 54.7 |

| | | | | |
|---|---|---|---|---|
| Pterin53 | 1.6 | 2.6 | 3.8 | 2.67 |
| Pterin54 | 14.3 | 16.1 | 16.3 | 15.57 |
| Pterin55 | 1.4 | 1.7 | 3.4 | 2.17 |
| 2-Nitroethanol | -4.2 | -4.3 | -4 | -4.17 |
| 2-Nitropropanol | -3.2 | -2.9 | -2.5 | -2.87 |
| 3-Nitropropionate | -5.3 | -5.1 | -5 | -5.13 |
| 3-Nitrooxypropanol | -5.5 | -5.4 | -5.2 | -5.37 |
| Bromoform | 0.2 | 1.2 | 2.6 | 1.33 |
| COB5 | 3.1 | 3.3 | 5.3 | 3.90 |
| COB6 | 4.3 | 6.1 | 6.2 | 5.53 |
| OCB7 | 12.1 | 12.4 | 12.4 | 12.30 |
| COB8 | 8.9 | 10.9 | 10.9 | 10.23 |
| COB9 | 13.4 | 14.1 | 14.6 | 14.03 |

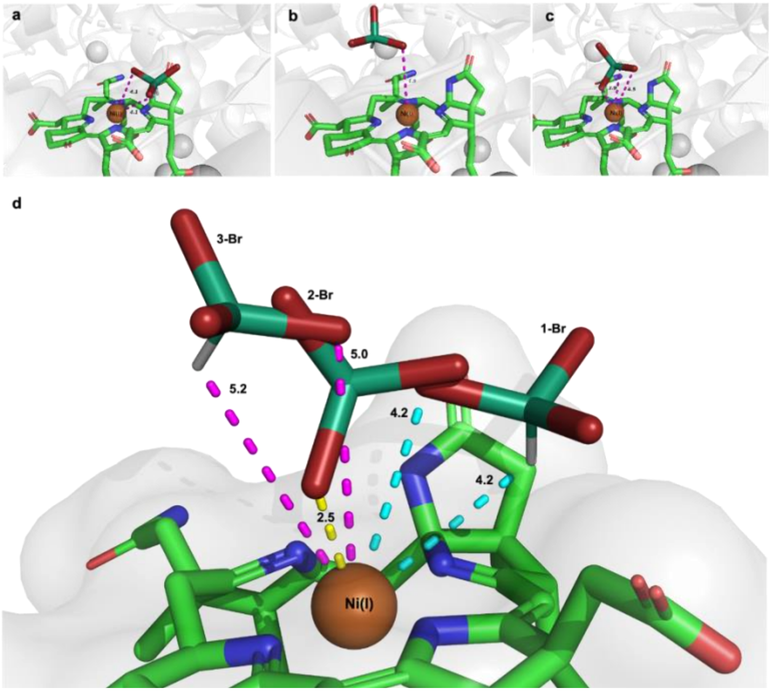


**Figure S2**. Illustration of all three non-superimposing poses of Bromoform molecule in the active site groove of MCR enzyme.

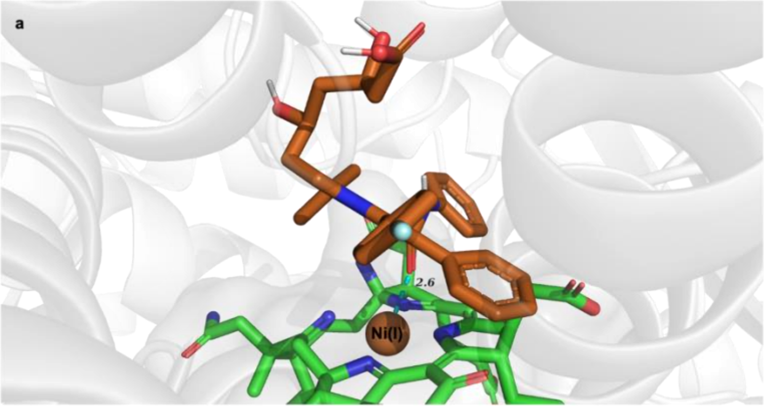


**Figure S3**. Illustration of the single pose of Atorvastatin molecule in the active site groove of MCR enzyme.

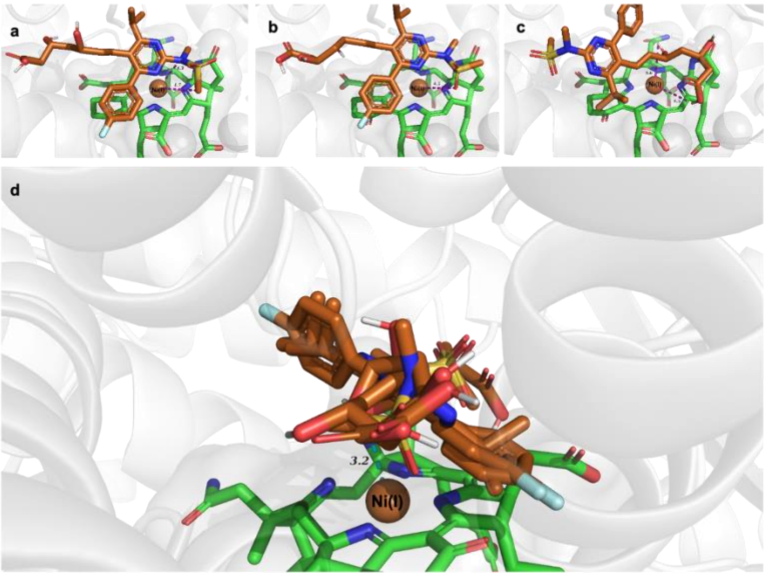


**Figure S4**. Illustration of all three poses of Rosuvastatin molecule in the active site groove of MCR enzyme.

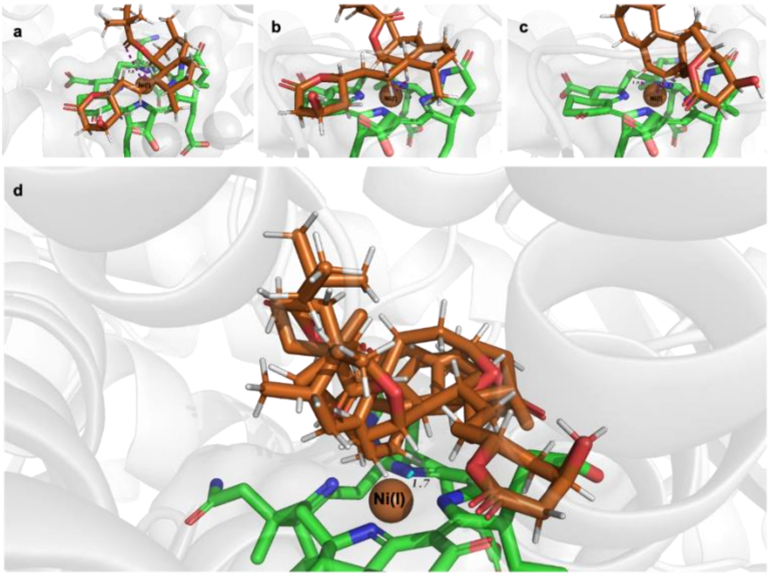


**Figure S5**. Illustration of all three poses of Simvastatin molecule in the active site groove of MCR enzyme.

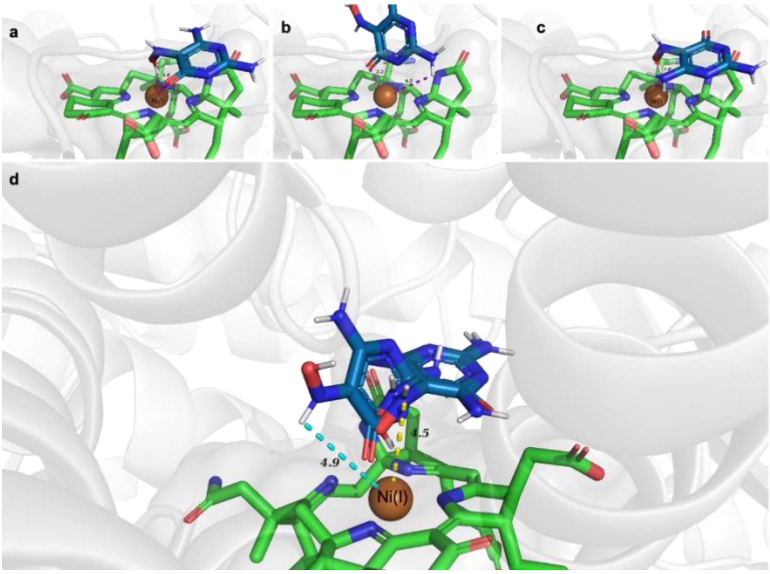


**Figure S6**. Illustration of all three poses of Pterin B53 (PT53) molecule in the active site groove of MCR enzyme.

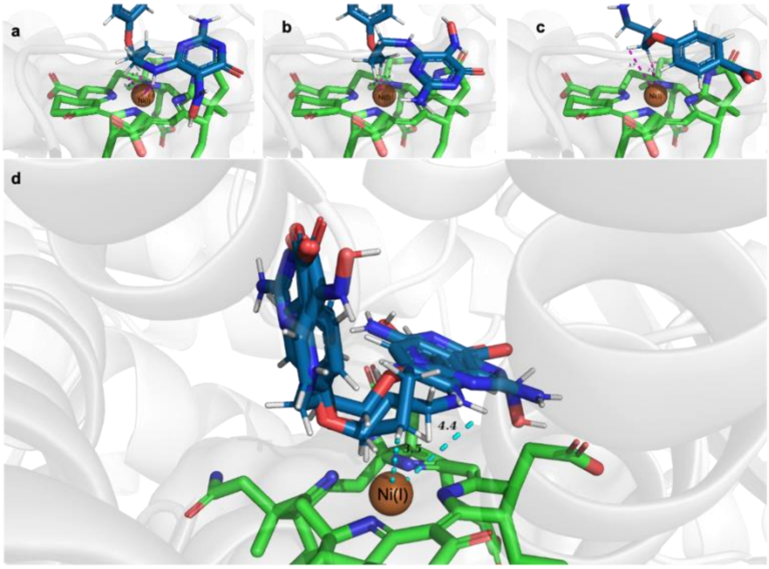


**Figure S7**. Illustration of all three poses of Pterin B54 (PT54) molecule in the active site groove of MCR enzyme.

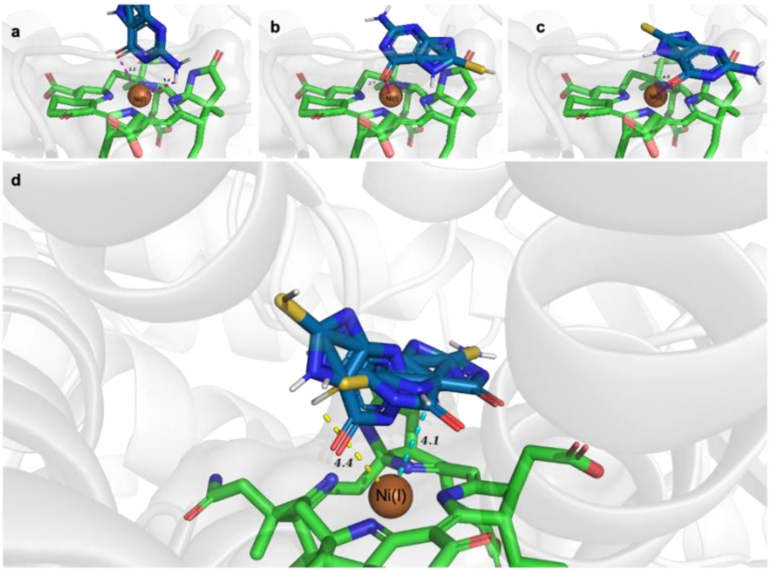


**Figure S8**. Illustration of all three poses of Pterin B55 (PT55) molecule in the active site groove of MCR enzyme.

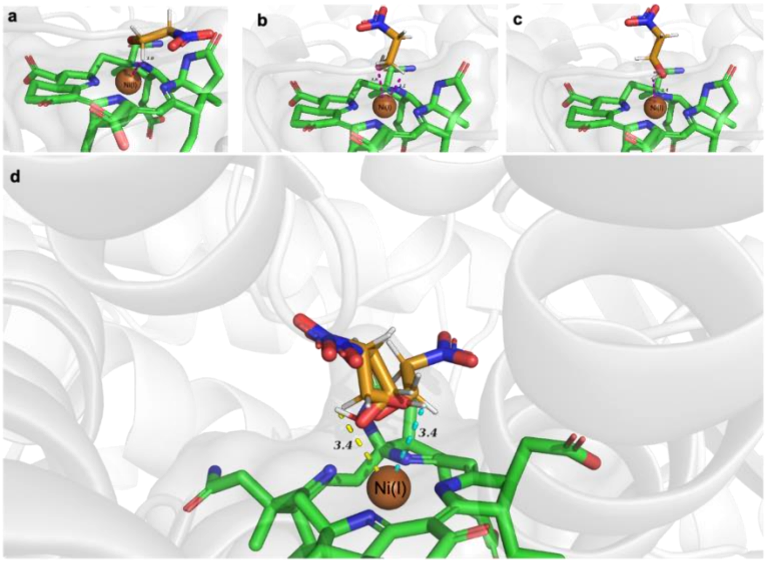


**Figure S9**. Illustration of all three poses of 2-nitroethanol molecule in the active site groove of MCR enzyme.

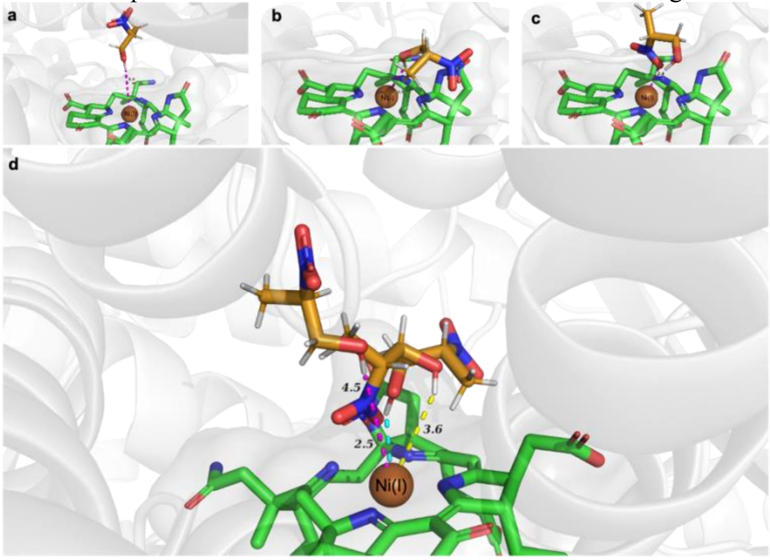


**Figure S10**. Illustration of all three poses of 2-nitropropanol molecule in the active site groove of MCR enzyme.

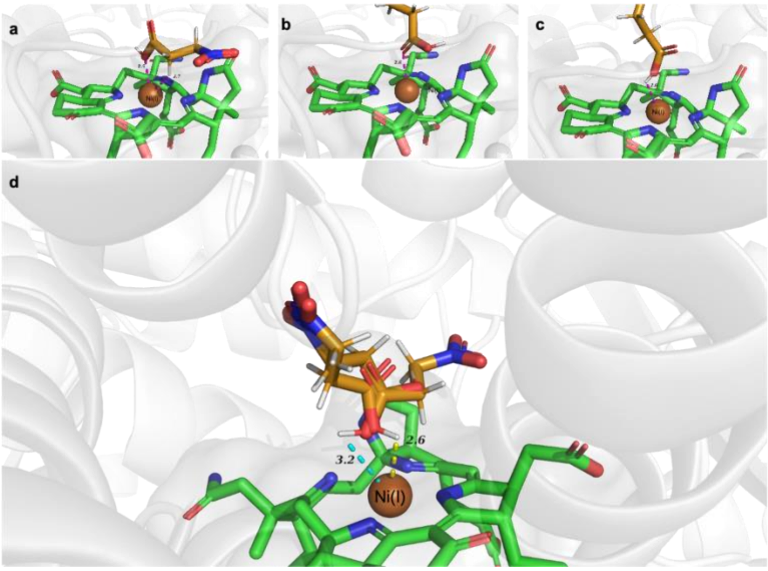


**Figure S11**. Illustration of all three poses of 3-nitropropionate molecule in the active site groove of MCR enzyme.

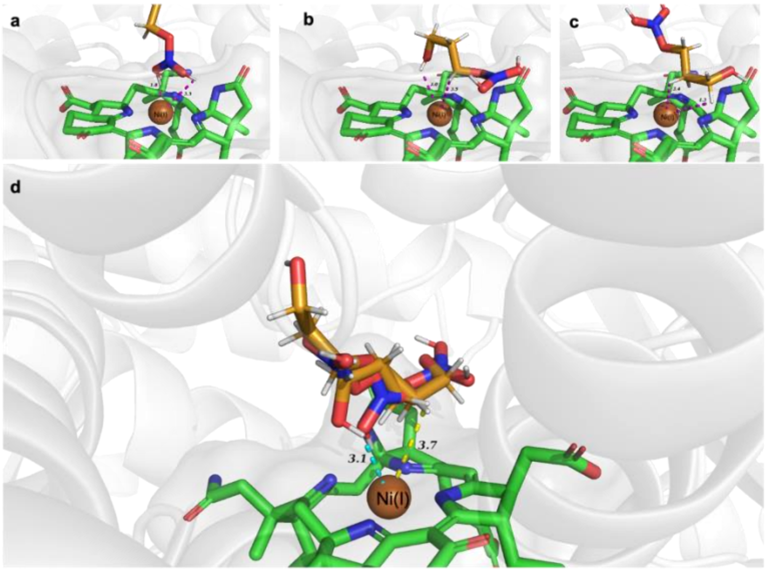


**Figure S12**. Illustration of all three poses of 3-nitrooxypropanol (3-NOP) molecule in the active site groove of MCR enzyme.

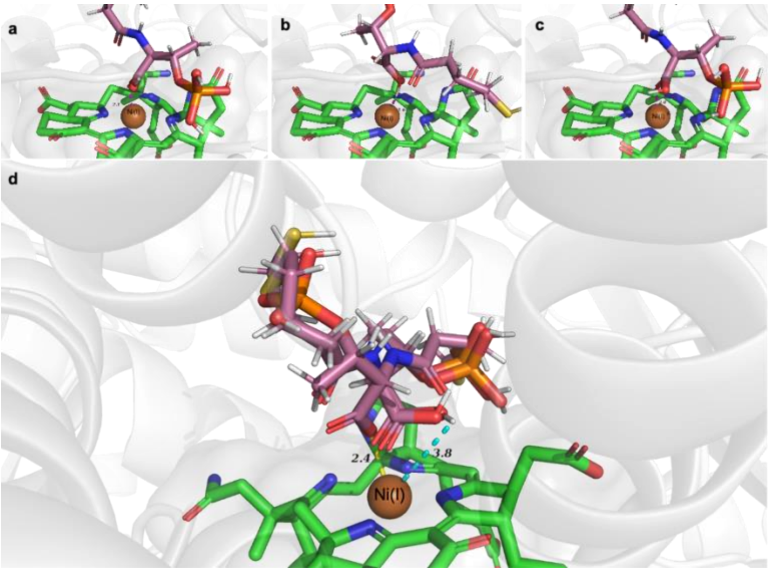


**Figure S13**. Illustration of all three poses of coenzyme B analogue (CoB5) molecule in the active site groove of MCR enzyme.

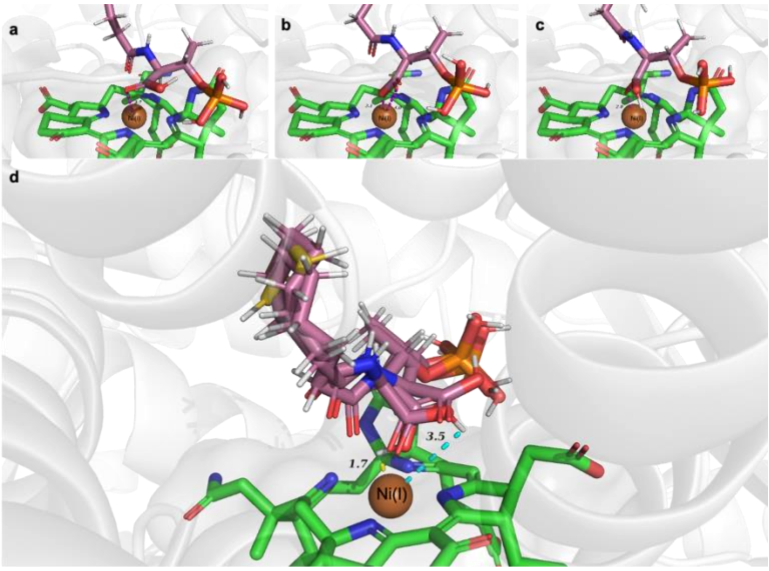


**Figure S14**: Illustration of all three poses of coenzyme B analogue (CoB6) molecule in the active site groove of MCR enzyme.

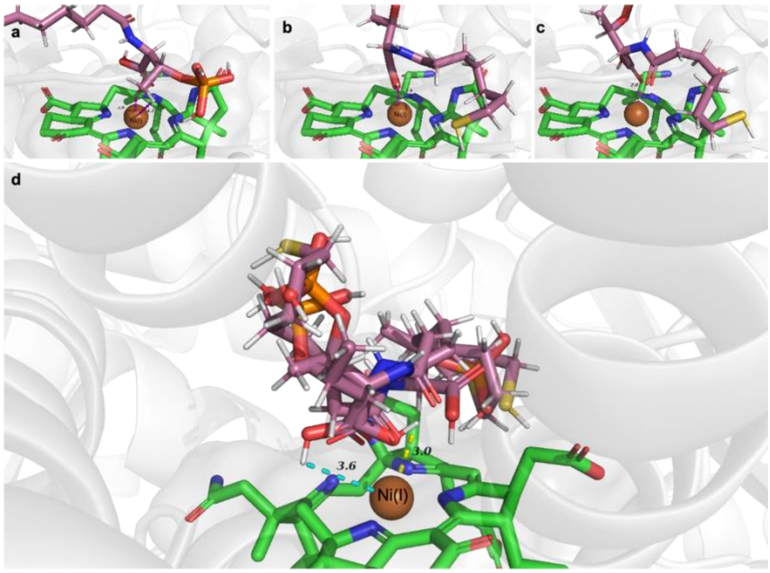


**Figure S15**: Illustration of all three poses of coenzyme B molecule (CoB or CoB7) molecule in the active site groove of MCR enzyme.

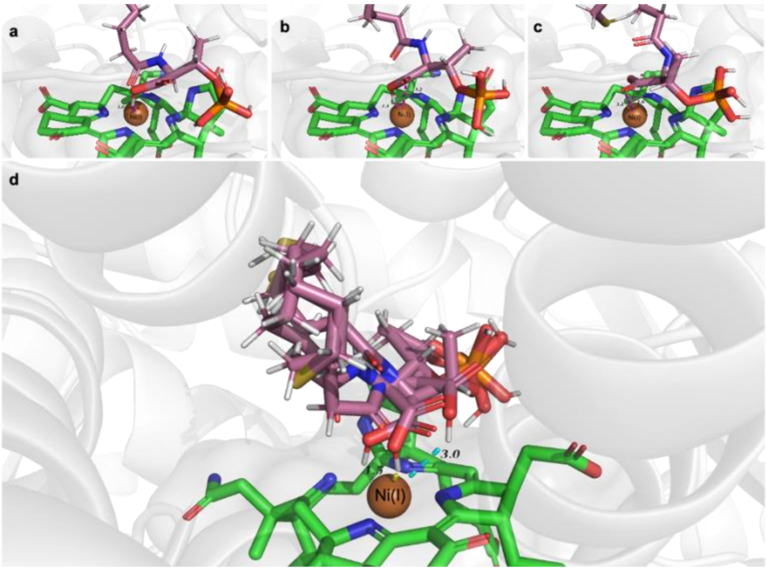


**Figure S16**: Illustration of all three poses of coenzyme B analogue (CoB8) molecule in the active site groove of MCR enzyme.

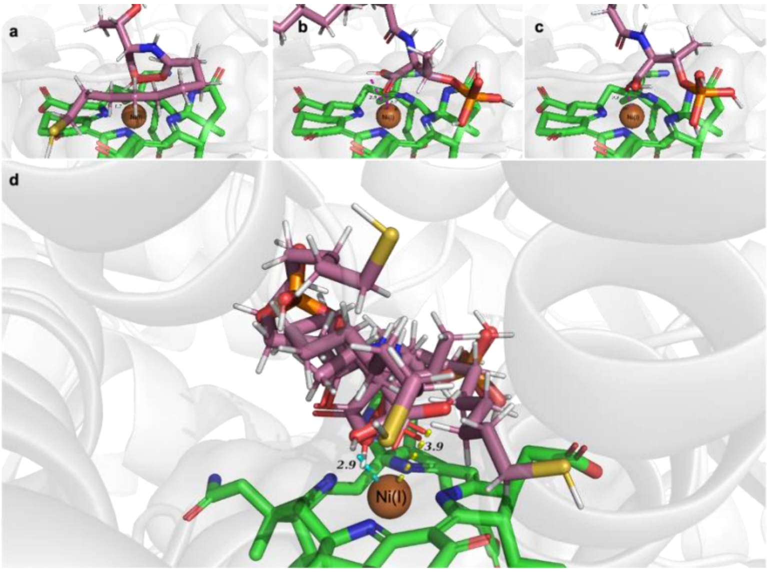


**Figure S17**: Illustration of all three poses of coenzyme B analogue (CoB9) molecule in the active site groove of MCR enzyme.

### Molecular Dynamics Simulations

Molecular Dynamics (MD) simulations were conducted utilizing the GROMACS molecular simulation platform[1,2] to systematically elucidate the interactions among MCR-cofactor $F_{430}$-anti methanogen ternary complexes. Protein was modelled using CHARMM36 force field while the cofactor was parameterized using ATB webserver.[3] Periodic boundary conditions were imposed in each dimension. Simulation box charge neutrality was achieved by adding $Na^+$ or $Cl^-$ ions, to avoid artifacts in the calculation of long-range electrostatics using Particle Mesh Ewald (PME).[4] We followed serial equilibration protocol- where initially, MCR- cofactor $F_{430}$ binary complex was equilibrated in a dodecahedral box of TIP3P[5] water at 300 K. This resulted in cofactor $F_{430}$ moving out of the solvation box and Ni(I) moved into the bulk solvent. We controlled the undesired movement of cofactor $F_{430}$ by using cubic water box, with 2.8 times increase in number of solvent molecules. Nevertheless, the tendency of Ni(I) to behave as a solvent ion continued to pose difficulty in modeling MCR-cofactor $F_{430}$ complex. Each molecular system was subjected to an energy minimization protocol using the steepest descent method.[6] This was followed by a 100-ps equilibration phase in the canonical ensemble at 300 K, with restraint force applied solely to the heavy atoms of the protein structure. The NVT stabilization phase was succeeded by an isobaric-isothermal (NPT) equilibration, rigorously maintained at a pressure of 1 bar and a temperature of 300 K, employing the Parrinello-Rahman barostat.[7] We attempted to add anti-methanogen (inhibitor) at the active site of the MCR, but it could not be handled as the system was not in well equilibrated state.

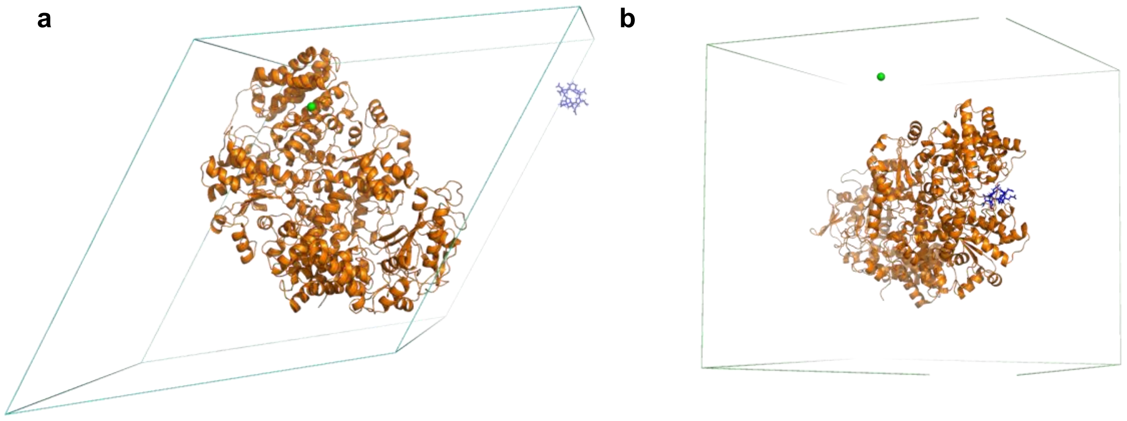


**Figure S18**: **a.** Dodecahedron simulation box with MCR (orange)-cofactor $F_{430}$. The cofactor F430 (shown in blue licorice) moved out of the simulation box as the simulation proceeded. Water molecules are omitted in the illustration for clarity. Ni(I) is represented by green bead. **b.** Cubic simulation box with MCR (orange)-cofactor $F_{430}$. The cofactor $F_{430}$ (shown in blue licorice) stays within the active site, but the Ni(I) ion moved towards bulk solvent as the simulation proceeded.

**Table 2**: List of BMDB and MCDB metabolites that are nearest and farthest to the known 16 inhibitors. BM- BMDB, MC- MCDB, N-Near molecules, F- Far molecules.

| Molecule | SMILES String |
|---|---|
| MCDB Near 1 | CCC(O)C(=O)O |
| MCDB Near 2 | CCCC(=O)O |
| BMDB Near 1 | O=C(O)CCCC(=O)C(=O)O |
| BMDB Near 2 | NCCCN |
| BMDB Near 3 | O=C(O)CC1=CC=C(O)C=C1 |
| MCDB Near 3 | CSCC(N)C(=O)O |
| BMDB Near 4 | O=C(O)CCCCC(=O)C(=O)O |
| MCDB Far 1 | C1=NC=NC2=C1N=C(NC2=O)C3CC(CO)O3 |
| MCDB Far 2 | C(C(=O)O)C(=C/C(=O)O)\C(=O)O |
| MCDB Far 3 | NCCO |
| BMDB Far 1 | CC1=CC2=C(C=C1)C=CC3=C4C=CC5=CC=CC=CC=CC(C4)=C3C(C2)=C5C=O |
| BMDB Far 2 | C(CC(=O)O)CN |
| BMDB Far 3 | C[C@]12CCC(=O)C=C1CCC1C2[C@@H](O)C[C@@]2(C=O)C1CC[C@@H]2C(=O)CO |
| BMDB Far 4 | NC1=NC(=O)C2=NC(CNC3=CC=C(C(=O)N[C@@H](CCC(=O)O)C(=O)O)C=C3)=CN=C2N1 |
| BMDB Far 5 | CCO |
| BMDB Far 6 | C=CC1=C(C)C2=CC3=C(C=C)C(C)=C(C=C4N=C(C=C5NC(=CC1=N2)C(C)=C5CCC(=O)O)C(CCC(=O)O)=C4C)N3 |

**Table 3:** Haddock Score of near and far metabolites after docking.

| Molecule | HADDOCK Score | Cluster Size | RMSD | Van der Waals Energy | Electrostatic Energy | Desolvation Energy | Restraints Violation Energy | Buried Surface Area | Z-Score |
|---|---|---|---|---|---|---|---|---|---|
| BMF1 | -18.7 | 19 | 0.1 | -11.5 | -28.4 | -5.3 | 9 | 273.5 | -1.6 |

| | | | | | | | | | |
|---|---|---|---|---|---|---|---|---|---|
| BMF2 | -18.8 | 9 | 0.1 | -8.8 | -68.5 | -4.1 | 10.2 | 253 | -1.1 |
| BMN1 | -13.7 | 12 | 0.1 | -7.2 | 0 | -7.6 | 10.6 | 259.1 | -1.6 |
| BMN2 | -18.6 | 5 | 0.1 | -14.1 | 0.5 | -5.7 | 11.4 | 281.4 | -1.4 |
| MCF1 | -12.7 | 42 | 0 | -8.7 | -3.7 | -4.5 | 7.9 | 223.5 | -1.4 |
| MCF2 | -15.8 | 31 | 0.1 | -7.2 | -80.3 | -1.4 | 8 | 195.2 | -1 |
| MCF3 | -15.9 | 6 | 0.1 | -9.2 | 0 | -7.5 | 8 | 257.1 | -1.1 |
| MCN1 | -9.4 | 10 | 0.1 | -5.5 | -1.4 | -4.6 | 8.6 | 190.3 | -0.9 |
| MCN2 | -12.5 | 28 | 0 | -7.7 | 0.2 | -6.8 | 20.1 | 247.4 | -1.1 |
| MCN3 | -15.4 | 16 | 0.1 | -10.1 | 0 | -6.2 | 8.9 | 266.8 | -1.7 |

**ML-based inhibitor search using graph neural network (GNN)**

A graph neural network (GNN) architecture was designed for clustering of known inhibitor molecules reported in literature against two unique databases - Milk Composition Database (MCDB) and Bovine Metabolome Database (BMDB), containing 2,360 and 51,682 entries, respectively. The significance for clustering was to explore for similarities in molecular fragments within these two ruminant databases against the known sixteen selected inhibitors of MCR enzyme. The structural information of metabolites was downloaded in Structure-Data File (SDF) format and further processed to obtain canonical Simplified Molecular Input Line Entry System (SMILES) representation using RDkit. Below are the smiles information used as input for the GNN.

```
#halo: Halogens
# coen: Coenzyme-B analogs
# nitro: Nitro-ol/esters
# stn: Statins
# ptrn: Pterins


# Format: Category|Generic Name|IUPAC Name|SMILES

halo|Bromoform|1,1,1-tribromomethane|BrC(Br)Br

coen|COB5|N-5-mercaptopentanoylthreonine
phosphate|P(=O)(O)(O)O[C@@H]([C@H](NC(CCCCS)=O)C(=O)O)C
coen|COB6|N-6-mercaptohexanoylthreonine
phosphate|P(=O)(O)(O)O[C@@H]([C@H](NC(CCCCCS)=O)C(=O)O)C
coen|COB7|N-7-
mercaptoheptanoylthreoninephosphate|P(=O)(O)(O)O[C@@H]([C@H](NC(CCCCCCS)=O)C(=O)O)C
coen|COB8|N-8-mercaptooctanoylthreonine
phosphate|P(=O)(O)(O)O[C@@H]([C@H](NC(CCCCCCCS)=O)C(=O)O)C
coen|COB9|N-9-mercaptononanoylthreonine
phosphate|P(=O)(O)(O)O[C@@H]([C@H](NC(CCCCCCCCS)=O)C(=O)O)C

nitro|2-Nitroethanol|2-Nitroethanol|[N+](=O)([O-])CCO
nitro|2-Nitropropanol|2-Nitropropanol|[N+](=O)([O-])C(CO)C
nitro|3-Nitropropionate|3-Nitropropionate|[N+](=O)([O-])CCC(=O)[O-]
nitro|3-Nitrooxypropanol [3NOP]|3-Nitrooxypropanol|[N+](=O)([O-])OCCCO
```

stn|Atorvastatin|(3R,5R)-7-[2-(4-fluorophenyl)-3-phenyl-4-(phenylcarbamoyl)-5-propan-2-ylpyrrol-1-yl]-3,5-dihydroxyheptanoic acid|FC1=CC=C(C=C1)C=1N(C(=C(C1C1=CC=CC=C1)C(NC1=CC=CC=C1)=O)C(C)C)CC[C@H](C[C@H](CC(=O)O)O)O
stn|Rosuvastatin|(E,3R,5S)-7-[4-(4-fluorophenyl)-2-[methyl(methylsulfonyl)amino]-6-propan-2-ylpyrimidin-5-yl]-3,5-dihydroxyhept-6-enoic acid|FC1=CC=C(C=C1)C1=NC(=NC(=C1/C=C/[C@H](C[C@H](CC(=O)O)O)O)C(C)C)N(S(=O)(=O)C)C
stn|Simvastatin|[(1S,3R,7S,8S,8aR)-8-[2-[(2R,4R)-4-hydroxy-6-oxooxan-2-yl]ethyl]-3,7-dimethyl-1,2,3,7,8,8a-hexahydronaphthalen-1-yl] 2,2-dimethylbutanoate|CC(C(=O)O[C@H]1C[C@H](C=C2C=C[C@@H]([C@@H]([C@@H]12)CC[C@H]1OC(C[C@@H](C1)O)=O)C)C)(CC)C

ptrn|Pterin B53|2,6-diamino-5-nitrosopyrimidin-4(3H)-one|NC1=NC(=C(C(N1)=O)N=O)N
ptrn|Pterin B54|4-{3-[(2-amino-5-nitroso-6-oxo-1,6-dihydropyrimidin-4-yl)amino]propoxy}benzoic acid|NC=1NC(C(=C(N1)NCCCOC1=CC=C(C(=O)O)C=C1)N=O)=O
ptrn|Pterin B55|2-amino-8-sulfanyl-1,9-dihydro-6H-purin-6-one|NC=1NC(C=2N=C(NC2N1)S)=O
